# Identification of the first endogenous *Ophiovirus* sequence

**DOI:** 10.1101/235044

**Authors:** Soledad Marsile-Medun, Humberto Julio Debat, Robert James Gifford

## Abstract

Endogenous viral elements (EVEs) are sequences in eukaryotic genomes that are derived from the ancestral integration of viral sequences into germline cells. Ophioviruses (family *Ophioviridae*) are a recently established family of viruses that infects plants. In this report, we describe the first example of an EVE derived from an ophiovirus, in the genome of eelgrass (*Zostera marina*). These findings extend the host range of ophioviruses to include seagrasses of the family *Zosteraceae*, and provide a potential time calibration for the evolution of the *Ophioviridae* family.

## Introduction

Ophioviruses (family *Ophioviridae*) are a recently established family of viruses that infects plants, causing economically important diseases [1, 2]. Only one genus (*Ophiovirus*) is currently recognized, containing seven species (Table 1). All ophioviruses are characterized by non-enveloped nucleocapsids that have helical symmetry and are highly filamentous. The negative-stranded RNA linear genome contains 3 or 4 segments coding for up to seven proteins. The first segment contains two ORFs, one encoding a 22–25K protein, and a second encoding the viral RNA polymerase. The second segment encodes the cell-to-cell movement protein (MP), while the coat protein (CP) is encoded by the third. A fourth segment has been reported in ophioviruses infecting lettuce (*Lactuca sativa*), which encodes putative proteins of unknown function [1].

**Table 1.** *Ophiovirus* reference sequences

| **Virus name** | **Abbreviation** | **Host** | **Accession** |
| --- | --- | --- | --- |
| Mirafiori lettuce big-vein virus | MLBvV | Lettuce | AAU12876 |
| Tulip mild mottle mosaic virus | TMmV | Tulip | AAT08133 |
| Lettuce ring necrosis virus | LRNV | Lettuce | YP_053239 |
| Freesia sneak virus | FSnV | Freesia | ABI33222 |
| Ranunculus white mottle virus | RWMV | Ranunculus | AAT08132 |
| Blueberry mosaic associated virus | BIMaV | Blueberry | APM86635 |
| Citrus psorosis virus | CPsV | Citrus | YP_089664 |

Endogenous viral elements (EVEs) are sequences in eukaryotic genomes that are derived from the ancestral integration of viral sequences into germline cells [3]. EVEs can provide unique retrospective information about the long-term coevolutionary history of viruses and their hosts [4, 5]. Here, we describe the first example of an EVE derived from an ophiovirus, in the genome of eelgrass (*Zostera marina*).

## Results

We screened genome assemblies of 142 plant species (**Table S1**) for sequences related to ophiovirus proteins. We identified only one statistically significant match, in the recently published genome assembly of eelgrass (*Zostera marina*) [6]. This sequence – hereafter referred to as *Zostera marina* endogenous ophioviral element (OphVe-ZosMar) spanned 567 nucleotides. When virtually translated, this sequence shared ~34–38% amino acid (aa) identity with the coat protein (CP) of known ophioviuses (**Figure 1**). The putative protein coding sequence of OphVe-ZosMar produced numerous, highly significant hits to ophioviruses when used to search GenBank, and no hits to sequences derived from other species.

**Figure 1.**
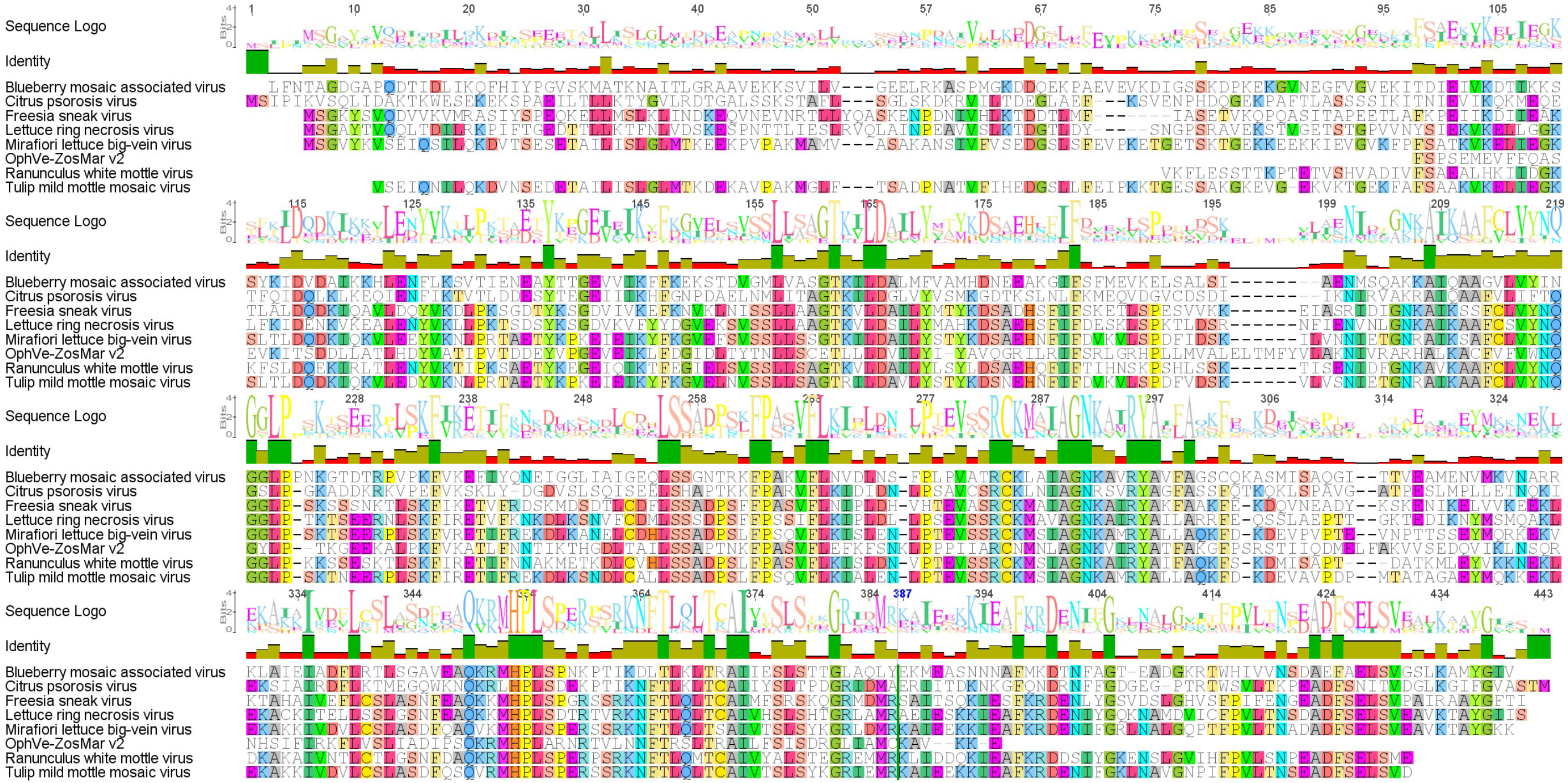
Alignment of ophiovirus coat protein (CP) sequences with the predicted polypeptide sequence encoded by *Zostera marina* endogenous ophioviral element (OphVe-ZosMar). Residues are coloured according to amino acid properties.

The OphVe-ZosMar element was identified in a large scaffold (accession # LFYR01000112.1: positions 103766-104332), and was flanked on either side by DNA sequences that exhibited no statistically significant similarity to any other viral or plant sequences in GenBank (**Figure S1**). However, regions flanking OphVe-ZosMar present protein domains typically associated with retrotransposons, indicating a plausible pathway for ancestral genome integration involving capture of viral mRNA by retrotransposable elements. Several RNA libraries of *Z. marina* have been published, and these contain OphVe-ZosMar derived reads, which supports the expression of this element (and the adjacent TE region).

Furthermore, neither the OphVe-ZosMar element nor it’s flanking sequences could be identified in the published genome assembly of *Zoster muelleri* [7], a related seagrass species. Assuming that the OphVe-ZosMar element is genuinely incorporated into the *Z.marina* genome (i.e. it does not reflect an artifact introduced through contamination), and is genuinely absent from the *Z.muelleri* genome, this would imply that the germline integration event that created OphVe-ZosMar occurred after these species diverged an estimated ~10-20 million years ago (MYA) [8, 9]. A sequence derived from a distantly related element (or a fragment of an related extant RNA virus) was identified in an RNA library of *Zostera noltei* (GenBank: HACV01019525.1). This fragment could potentially indicate the presence of an orthologous insert in a second seagrass species, but amino acid identity with OphVe-ZosMar was relatively low (**Figure S2**), suggesting that they are derived from a distinct virus or germline incorporation event.

We used maximum likelihood to infer the phylogenetic relationships between known ophioviruses and the OphVe-ZosMar. As shown in **Figure 2**, the phylogeny discloses two well-supported subgroups within the *Ophioviridae*, one that contains the ophioviruses infecting citrus and blueberry (**group 1**), and another containing all other ophioviruses (**group 2**). In midpoint rooted trees, the OphVe-ZosMar element groups outside both of these clades. However, the branch length separating OphVe-ZosMar from these two groups of exogenous ophioviruses was not much greater than that separating the two groups from one another.

**Figure 2.**
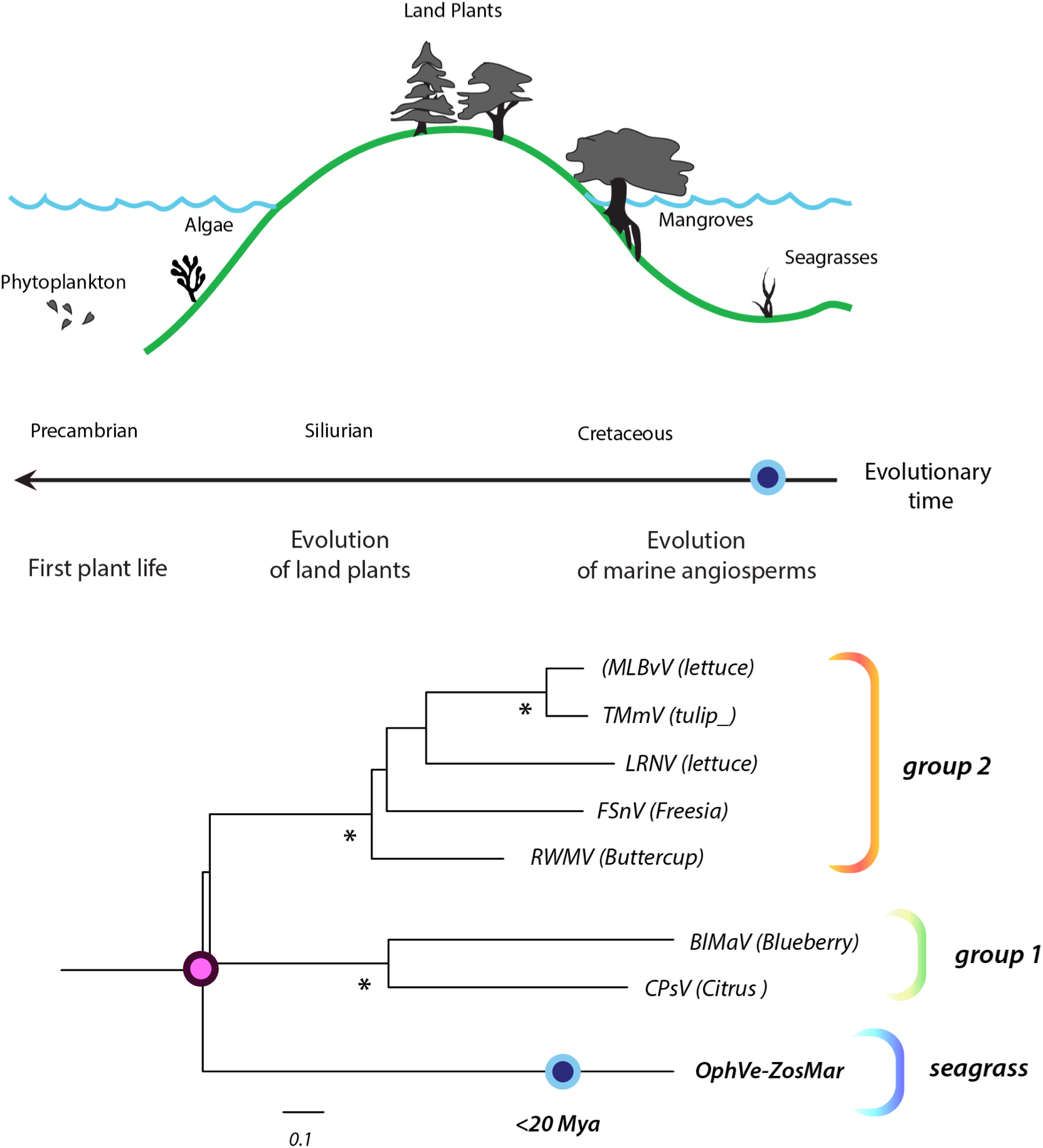
The bottom panel shows a bootstrapped maximum likelihood phylogeny with the inferred evolutionary relationships between *Zostera marina* endogenous ophioviral element (OphVe-ZosMar) and representative ophioviruses. The tree was inferred from the polypeptide alignment shown in **Figure 1**, and is midpoint rooted for display purposes. Asterisks indicate nodes with >95% bootstrap support. The scale bar shows phylogenetic distance in substitutions per site. Accession numbers of ophiovirus taxa shown here are given in **Table 1**. The top panel juxtaposes the phylogeny of ophioviruses against the evolutionary history of angiosperms, showing how seagrasses evolved from land plants. Our investigation of other seagrass species, suggests OphVe-ZoMar may have inserted <20 million years ago (Mya), as indicated by the blue dot. To the extent that seagrass viruses are isolated from viruses infected plants on land, the pink dot should correspond approximately to the point at which these populations became isolated. Adding further time-points (e.g. through identification of other seagrass ophioviruses or endogenous viral elements) would allow more confident calibration of ophiovirus evolution.

## Discussion

Progress in whole genome sequencing has led to the identification of numerous, diverse EVE sequences in the eukaryotic species. Many of these EVEs are clearly derived from well-recognized virus families, whereas others are only distantly related to contemporary viruses. Furthermore, although EVEs derived from a diverse range of virus families have been, there are still many virus families for which no EVEs have been identified.

In this paper we describe the identification of *Zostera marina* endogenous ophioviral element (OphVe-ZosMar) – the first reported example of an EVE derived from an ophiovirus. This sequence, which was derived from the segment of the ophiovirus genome that codes for the viral coat protein (CP), was identified in the recently sequenced genome of eelgrass (*Zostera marina*) [6]. The DNA sequences flanking OphVe-ZosMar did not disclose significant similarity to genome sequences identified in other plants, and were also found to be moderately repetitive. For this reason, we could not completely rule out the unlikely possibility that the presence of OphVe-ZosMar in a large contig reflected an artifact associated with contamination and/or misassembly during the production of the *Z. marina* genome. Future studies of seagrass genomes should allow the presence of the OphVe-ZosMar element in *Z. marina* to be confirmed. Confirmation of that the OphVe-ZosMar element occurs in eelgrass would enable further investigations, in particular, it should allow the age of the element to be estimated, providing some insight into the timeline of ophiovirus evolution, about which nothing is currently known. In addition, since the OphVe-ZosMar element appears to encode an intact (or nearly intact) CP protein, and there is evidence from DNA libraries that this element is expressed, the possibility of conducting functional studies may also exist.

Seagrasses are one of several groups of angiosperms (flowering plants) that, having evolved on land, subsequently colonised the marine environment [10]. Terrestrial plants are thought to have originated during the Silurian period (~450 MYA), but it was not until ~130 million years ago that angiosperms evolved and invaded marine environments. Assuming that colonization of the marine environment isolated ophiovirus populations infecting seagrasses from those infecting their ancestors on land, this event could potentially be used – in combination with dating of OphVe-ZosMar – to calibrate the evolution of the ophiovirus family (**Figure 2**). Further characterization of EVEs in marine angiosperm genomes can potentially provide some insight into how this macroevolutionary shift impacted the distribution and diversity of plant viruses.

## Methods

### Genome screening and sequence analysis

Plant genome assemblies (**Table S1**) were downloaded from NCBI (www.ncbi.nlm.nih.gov/genome/). Screening was performed using the database-integrated genome-screening tool (available from http://giffordlabcvr.github.io/DIGS-tool/). ORFs were inferred by manual comparison of putative peptide sequences to those of closely related exogenous parvoviruses in the alignment editing software Se-Al [11]. The putative peptide sequences of *Ophiovirus-related* EVEs were aligned those of representative ophioviruses using MUSCLE [12] and PAL2NAL [13]. Phylogenies were reconstructed from this alignment, using maximum likelihood as implemented in RaxML [14], and the VT protein substitution model as selected using ProTest [15].

## Compliance with Ethical Standards

RJG was funded by the Medical Research Council of the United Kingdom (MC_UU_12014/12).

### Ethical approval

No humans or animals were involved in this study

